# Arginine versus Lysine: Molecular Determinants of Cation–π Interactions in Biomolecular Condensates

**DOI:** 10.1101/2025.10.01.679751

**Authors:** Lydia Armentia, Xabier López, David De Sancho

## Abstract

Biomolecular condensates formed by intrinsically disordered regions of proteins are primarily stabilised by specific types of amino acids, usually labelled “stickers” for their ability to crosslink distinct polypeptide chains into droplet-spanning networks. Both aromatic and positively charged residues can play this role, but they differ in their interaction types and relative strengths. An outstanding problem is the quantification of these differences, since they determine the fate of multicomponent mixtures. Here we focus on the interactions formed by aromatic residues with the two main positively charged stickers, Arg and Lys, whose experimental behaviour in condensates shows a clear hierarchy: Arg is consistently observed to promote phase separation more effectively than Lys. We use molecular dynamics simulations and alchemical transformations together with quantum chemical calculations to resolve the differences in their cation–*π* interaction strengths in different media. We find that, unlike the aromatic residues Phe and Tyr, Arg and Lys consistently maintain a hierarchy of interaction strengths across molecular environments, with Arg being favoured over Lys. While cation-*π* interactions are important, the primary factor underlying this difference is the higher dehydration penalty of Lys. By contrast, the identity of the aromatic partner that forms the strongest interaction with a cation depends on the dielectric environment. These results provide a clear molecular-level explanation for the distinct contributions of interactions between cationic and aromatic residues to condensate stability.

## INTRODUCTION

In recent years, our understanding of cellular organisation has been transformed by the coming to prominence of membraneless organelles, formed primarily by proteins and nucleic acids [1]. These organelles can form both in the cytoplasm, such as P-bodies and stress granules, and in the nucleus, as in the case of nuclear speckles, Cajal bodies, or the nucleolus. The idea that membraneless organelles may arise through liquid–liquid phase separation of their constituents has renewed interest in the phase behaviour of biomolecules.

Much of our understanding of condensate formation is mediated by theories of polymers [2]. The “stickers and spacers” model borrowed from the theory of associative polymers [3–6] has been particularly successful as a framework for interpreting experiments on condensates, both in ordered [7] and disordered proteins [5, 8]. In this model, different amino acid residues or groups thereof are assigned specific roles, either mediating the cross-links between interacting units (“stickers”) or providing the necessary flexibility and fluidity to the resulting network (“spacers”). In the case of intrinsically disordered proteins (IDPs) or regions (IDRs), experiments have resolved a specific grammar for phase separation [5]. Aromatic amino acid residues (Phe, Tyr) and positively charged (Lys, Arg) residues appear prominently in the sticker category. Polar residues, like Ser, and Gly play the role of spacers. While all amino acid residues can form hydrogen bonds, the distinct chemical nature of different stickers allows them to form distinct types of interactions, including π-stacking, cation-π or charge-charge, all of which are relevant in condensates [9].

An important question to understand these condensates more fully is what is the relative strength of the different stickers. Experimentally, this problem has been approached via ingenious sequence perturbations in phase-separating proteins [5, 10–15]. A key result is that interactions between Arg and Tyr govern condensation and that they are much stronger stickers than their counterparts Lys and Phe [5]. In fact, the number of Arg and Tyr residues alone can be predictive of the saturation concentration of a phase-separating protein [5], although contributions from other residue types are also important [16].

We have recently studied the molecular origin of the greater stickiness of Tyr relative to Phe in biomolecular condensates [17]. The difference between Tyr and Phe was partly unexpected considering that Phe is broadly regarded as more hydrophobic than Tyr [18, 19] and hydrophobicity is one of the key driving forces for phase separation [20, 21]. We employed molecular dynamics (MD) simulations of minimal model condensates [22] and quantum chemical calculations to demonstrate that the relative sticker strength is significantly influenced by the difference in transfer free energy, which depends on the environment [17]. Transferring stickers from water into the cores of folded proteins, with a very low dielectric constant [23], is more favourable for Phe. However, in the case of condensates —a highly solvated medium with intermediate dielectric [24, 25]— the balance tilts towards Tyr [17].

Here we turn our attention to the positively charged residues Arg and Lys. As with Phe and Tyr, one may naively assume that these two residues are interchangeable, given their shared ability to engage in cation-π interactions and similarly high pK_a_ values in model compounds (13.9 and 10.34, respectively [26]). However, experiments on FUS [5], Ddx4 [27] LAF-1 [13] or MCDs [12], where R→K mutants reduced or entirely suppressed phase separation, point to a greater sticker strength of Arg. The relative strengths of Arg and Lys as stickers have also been interrogated using computational approaches, including simulations with coarse-grained models [25] and atomistic force fields [28–30], and computational chemistry calculations [31]. A consensus is emerging that while both share their role as cations, Arg is the preferred interaction partner of aromatics like Phe and Tyr. A contributing factor to these differences is the pseudo-aromatic character of the guanidinium group, where the positive charge is delocalized [32]. This facilitates the dehydration of the guanidinium plane in Arg, promoting interactions with other residues [33]. Additionally, while bound to an aromatic residue, Arg can participate in hydrogen bonds with water or other amino acids [34, 35].

In this work, we use a combination of non-equilibrium alchemical transformations [36] and density functional theory (DFT) calculations to examine whether the context-dependent stickiness observed for Phe and Tyr applies also to the cation–π pairs formed with Arg and Lys. We find that Arg is consistently a stronger sticker than Lys, irrespective of the environment, while the dependency on the dielectric observed in our previous work determines whether Phe or Tyr are the most favourable partner to form cation-π pairs. Our results highlight that, in addition to the interaction strengths of Lys and Arg within the condensate, a key distinction between both residues lies in their differing free energies of transfer from water.

## RESULTS AND DISCUSSION

### Alchemical transformations reproduce the experimental hierarchy of sticker strengths

Transfer free energies of aminoacid residues from water into condensates are an important contribution to sticker strength in biomolecular condensates [17, 37]. Hence, we first calculate the relative transfer free energy (ΔΔG_Transfer_) into a model condensate for a peptide bearing either Lys or Arg. Direct simulations to estimate the values of ΔG_Transfer_ are very challenging. Alternatively, we use the thermodynamic cycle in Figure 1, where we are representing the molecule of interest (GGXGG) in both chemical states (X=K and X=R) submerged in two different solvents (water and a model condensate). In the diagram, the transfer process is represented by the vertical arrows and associated with the free energy 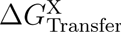. Because we are interested in the difference in transfer free energies between Arg and Lys, we can instead calculate the free energies ΔG_K→R_ in either water or the condensate associated with the horizontal branches, which involve an alchemical transformation. Because of the path-independence of free energy changes, we can write

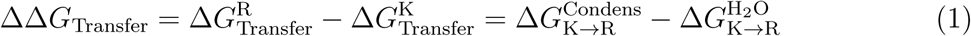

**FIG. 1.**
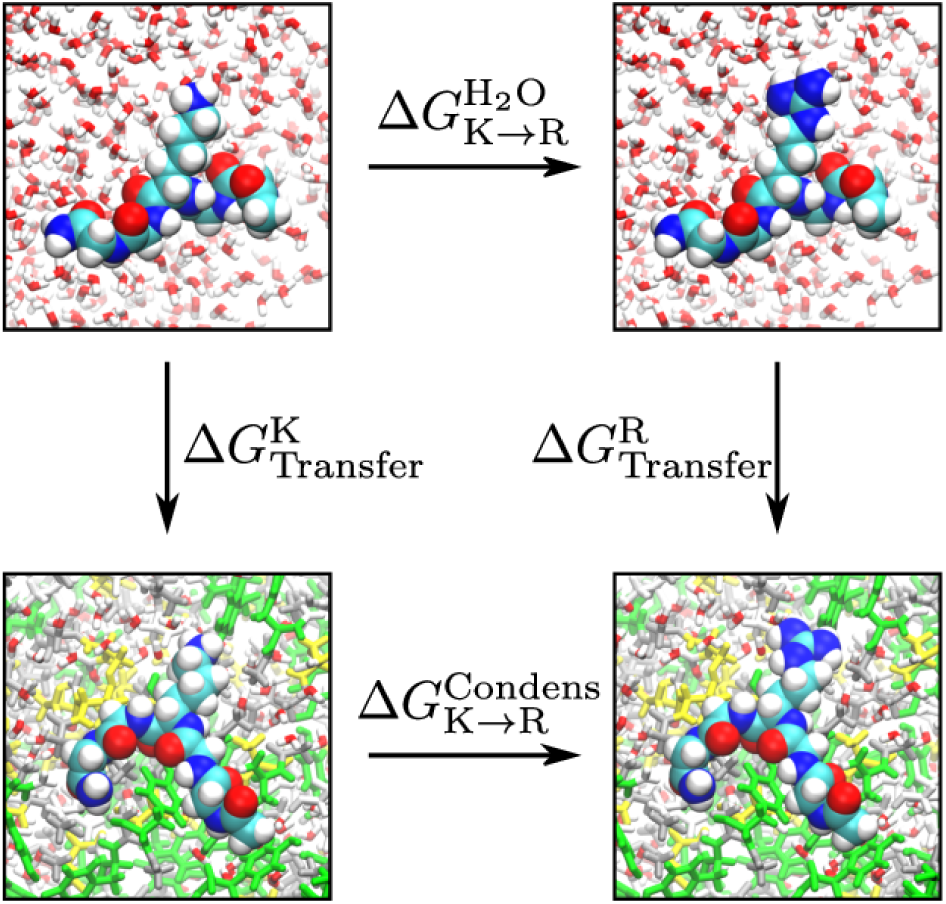
Thermodynamic cycle for the estimation of transfer free energy differences from alchemical MD simulations. The cycle involves the transfer of short peptides with sequence GGXGG from water to condensates (vertical branches) and the alchemical transformation from X=Lys to X=Arg (horizontal branches). In the condensates, different colours indicate different constituent residues (Gly: white; Ser: yellow; Tyr: green).

Hence, using the values for ΔG_K→R_ in water and in the condensate, which are easier to calculate, we can determine ΔΔG_Transfer_.

We have first run long equilibrium simulations of the GGXGG pentapeptide in the λ = 0 (X=K) and λ = 1 (X=R) states inside ternary peptide condensate slabs formed by capped Gly, Ser and either Tyr or Phe (termed GSY and GSF, respectively). Because the peptide is positively charged, we have run two different neutralization conditions to control for their influence in our results (see Methods). We show a representative snapshot of the GSY condensate in Figure 2A. The boxes take up to 500 ns to equilibrate, as inferred from the convergence of the solvent accessible surface area of the protein atoms (SASA) (see SI Figure S1). From that point, the condensate remains stable. In the case of the GSY condensate, the GGXGG peptide remains buried throughout most of the long simulation trajectories, although we see it escape in the box neutralised with just one Cl^−^ anion for the λ = 0 (see SI Figure S2B). We show the average density profiles for the equilibrated part of the simulation trajectories in physiological conditions in Figure 2B (see also SI Figure S2). In the case of the less protein-dense GSF condensate, the GGXGG peptide escapes from the condensate interior more readily and stays temporarily at the interface region (see SI Figure S3). These simulation results are consistent for both systems irrespective of the procedure used to neutralise the simulation box (see SI Figures S1-S3).

**FIG. 2.**
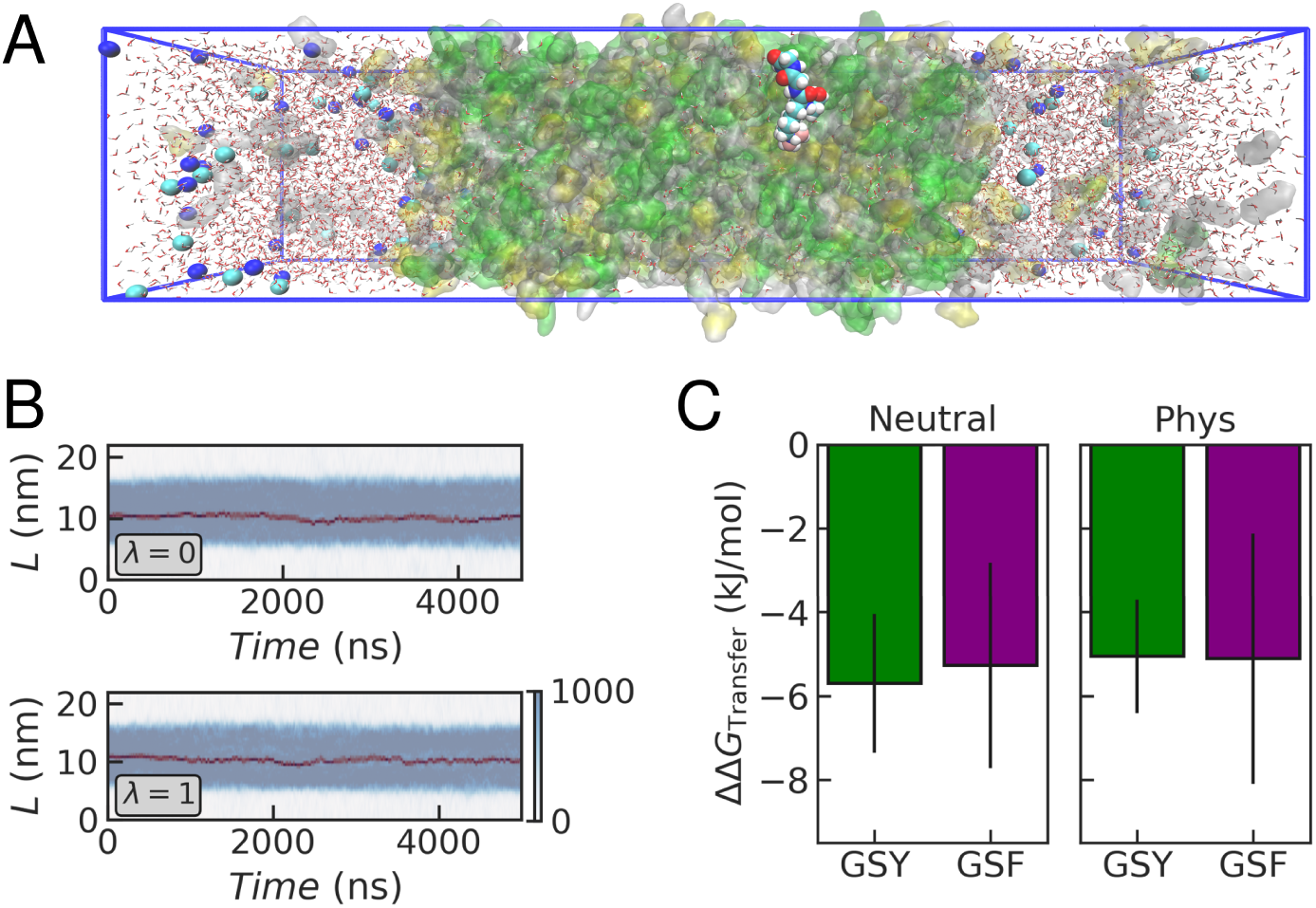
Classical MD simulations of the GGXGG peptide inserted in the peptide condensates. (A) Representative snapshot of GGXGG in the GSY slab at physiological salt. The condensate is shown as a transparent surface with residues shown in different colours (G: White, S: yellow, Y: green). The peptide and Na^+^ and Cl^−^ ions are shown as spheres. (B) Density profiles as a function of time for the end states *λ* = 0 (X=K) and *λ* = 1 (X=R) in the GSY condensate at physiological salt. The total protein density is shown in blue, and density of the K/R residue is shown in dark red. Density is expressed in mg/mL. (C) Estimates of the transfer free energy difference at neutral and physiological ionic strength in the GSY and GSF condensates.

We selected equally spaced snapshots from the long simulation runs, excluding the first 500 ns of the trajectories and any configuration where the peptide was not buried within the condensate. Then we ran hundreds of fast, non-equilibrium switches from both end states. Using the statistics of (∂H/∂λ) for all initial states, we estimate the values of 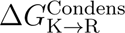 using the PMX workflow [36] (see Methods and SI Figure S4). Switches were also run for simulations of the peptide in both λ states in water matching the neutral or physiological conditions, in order to obtain 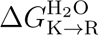. Combining both free energy estimates, we obtain ΔΔG_Transfer_ using Equation 1. We find that the value of ΔΔG_Transfer_ is negative in both the GSF and GSY condensates (see Figure 2C). This indicates that Arg has a stronger propensity to be part of the model peptide condensates than Lys. The results are independent of the neutralisation conditions. Hence, the emerging hierarchy of sticker strength based on our calculations is Arg-Tyr > Lys-Tyr and Arg-Phe > Lys-Phe. This result is consistent with a plethora of experimental and computational results that point to a higher stickiness in Arg [5, 12, 13, 25, 27, 29, 38].

Several interesting comparisons can be drawn between the current results and those obtained for Tyr and Phe in our previous work [17]. For this comparison, we consider the “neutral” conditions, as no ions were added in the F→Y transformations. Although Tyr was found to be more “sticky” than Phe based on the negative ΔΔG_Transfer_ of the F→Y transformation, the difference between aromatics was small (∼ 1.5 kJ/mol). For the K→R transformation, the magnitude of the differences in transfer free energy is much greater (∼ 5 ± 2 kJ/mol). This is consistent with the generally more disruptive effect of R→K mutations, often capable of completely suppressing phase separation [13, 27] and with the magnitude of free energy differences in potentials of mean force obtained from MD simulations [30, 39].

### Context independence of Arg and Lys transfer free energies

In our previous work, we reported that the hierarchy of sticker strengths between Tyr and Phe is context-dependent [17]. This effect is governed by the magnitude of the transfer free energy of the amino acid residue from water into a different environment. Tyr is favoured for polar media where it can form hydrogen bonds, such as the water-rich GSY and GSF condensates or acetone, whereas Phe is preferred in apolar solvents like cyclohexane or benzene. We found that the value of the dielectric constant ε of the media tracks this crossover. This is important because ε for condensates is around 40, roughly half that of water [17, 24, 25], and it is much lower in the hydrophobic cores of folded proteins [40].

In the case of Arg and Lys we have similarly interrogated the dependence of the relative transfer free energy on the target medium. We have performed the same type of alchemical transformation in multiple other solvents and estimated the value of ΔΔG_Transfer_ (see SI Figure S5). In this case, we find that irrespective of the solvent, ΔΔG_Transfer_ is always negative, and hence consistently favours Arg relative to Lys (see Figure 3A). Although there are quantitative differences across solvents, we cannot establish a systematic dependence of ΔΔG_Transfer_ on ε (see Figure 3B). Also, the numerical results for toluene, benzene and cyclohexane may be influenced by the difficulty of sampling the equilibrium distribution of peptide torsion angles in these solvents, which can affect the outcome of the non-equilibrium switching [41] and the potentially unrealistic background neutralizing effect of the particle mesh Ewald (PME) method [36]. In any case, the results of the alchemical transformations suggest a robust trend for a more favourable transfer free energy of Arg across diverse environments.

**FIG. 3.**
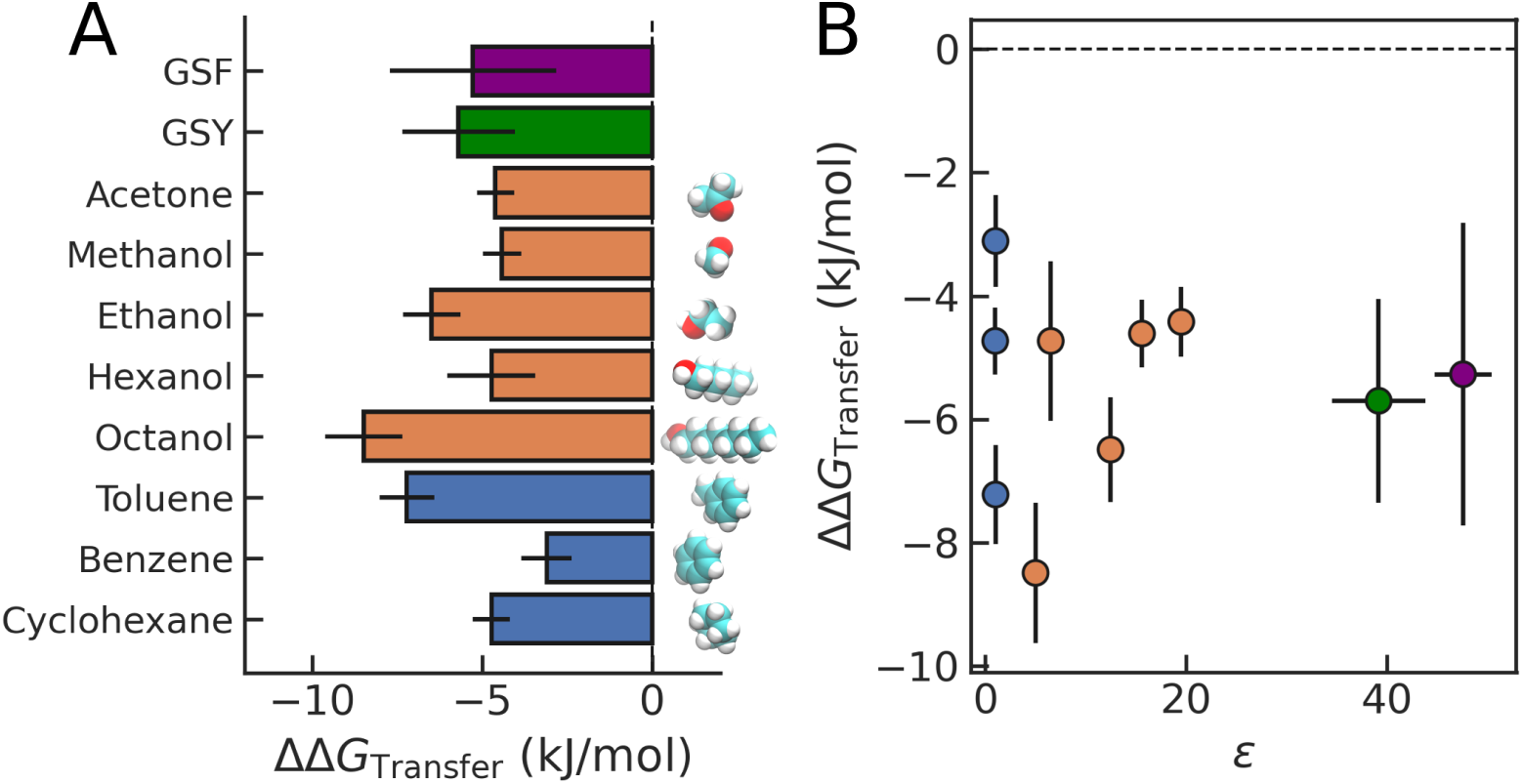
Transfer free energy differences from alchemical transformations. (A) Values of ΔΔ*G_T ransfer_* in different solvents. (B) Relationship between calculated dielectric constants and transfer free energy differences.

### Interaction modes of Lysine and Arginine in peptide condensates

We next investigate the interactions formed by Arg and Lys inside the peptide condensates. For this, we study the contacts formed by the sidechain of the alchemically transformed residue during the long equilibrium MD runs. We have inspected the values of inter-sidechain distances and the relative orientations of the Arg guanidinium group and the aromatic ring of either Tyr (in the GSY condensate) or Phe (in GSF), characterized by the angle θ between the vectors normal to each plane. Note that the latter can be calculated for both the GGKGG and the GGRGG peptide due to the dual topology approach used in our simulations. We find a clear propensity to form stacked arrangements between the Tyr/Phe and Arg sidechains (λ = 1), which appear at short distances and values of θ close to 0 (see Figure 4 A, C). In the case of Lysine (λ = 0), there is no orientational preference for the charged sidechain relative to the aromatic ring, as expected. This trend is consistent for both GSY and GSF condensates (see representative snapshots of stacked configurations in Figure 4 B, D and additional conformational analysis in Supporting Figures S6 and S7). This clear distinction in contact patterns between Arg and Lys supports a role for π-stacked interactions in the more favourable transfer free energy of Arg. The preference for π-stacking in the simulations is consistent with the abundance of this type of arrangement in experimental structures of proteins [34, 42] and the strongly attractive potentials of mean force reported for Arg–aromatic pairs using both fixed-charge and polarizable force fields [28, 30, 43]. Nevertheless, this type of geometry may arise due to its efficient packing of sidechains or the ability to fulfil other interactions simultaneously [34], rather than from an intrinsically stronger interaction.

**FIG. 4.**
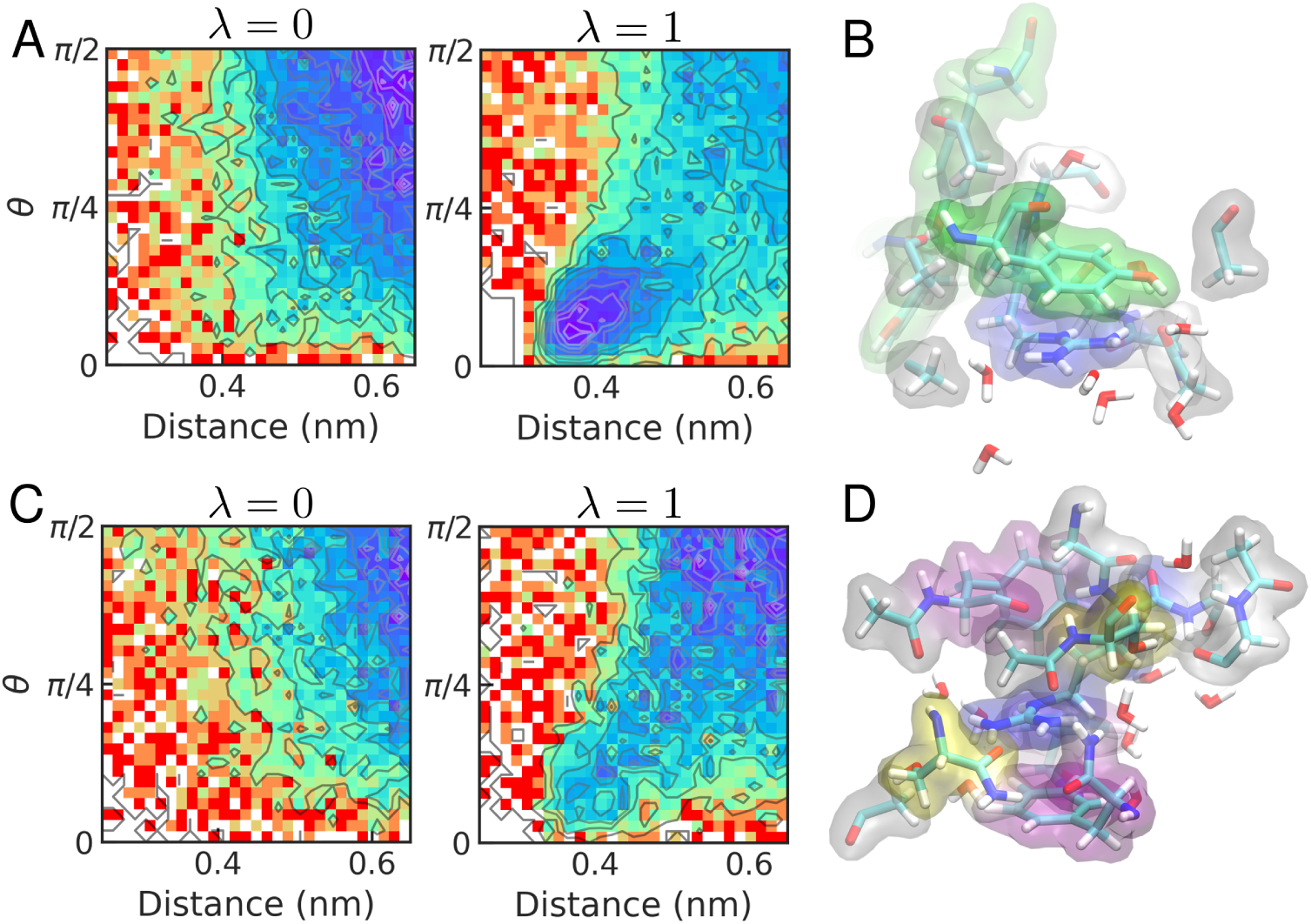
Conformational preferences of Lys and Arg in peptide condensates. (A) Density plots of the distance and angle *θ* formed between normal vectors to the aromatic ring and guanidinium plane of the GGXGG peptide in the GSY condensate. *λ* = 0 corresponds to X=K and *λ* = 1, to X=R. In the case of X=K, we use the dummy atoms in the dual topology to measure *θ*. (B) Representative snapshots of *π*-stacked arrangement of Tyr and Arg. (C-D) Same as A and B, respectively, for the GSF condensate.

A potential limitation of our results is that the cation-π interaction can only approximately be reproduced by classical force fields. This is due to the lack of a proper representation of the polarizing effect of the cation over the π-electrons [44–46]. Recently, different approaches have been proposed to overcome these limitations, including an empirical correction to the Lennard-Jones potential [47] or ad-hoc modifications in the charges for bound states [28]. Another possibility is to resort to polarizable models to study the hierarchy of interactions [30]. Alternatively, here we resort to quantum mechanical calculations using DFT.

### Contact energies from quantum chemical calculations

Our investigation of the interactions between positively charged and aromatic residues reveals an interplay of intrinsic forces and environmental influences. To gain insight into these effects, we performed DFT calculations considering the side chains of Lys, Arg, Phe, and Tyr (see Methods). Specifically, we evaluated the interaction energies (ΔE_Int_) of all possible charged-aromatic dimers in various configurations, taking into account environmental conditions of different polarities. To introduce environmental effects, two types of solvents were considered within a polarizable continuum model, as before [17]: (i) solvents of different polarity and (ii) water with a reduced dielectric constant, which may capture the effect of an increasingly restrained water solvent. Although this second solvent model may seem artificial, a dielectric decrement occurs both in electrolyte solutions [48] and when water is nanoconfined [49]. The latter scenario may not be far removed from that of biomolecular condensates, where water molecules diffuse within a polymer network with a mesh size in the range of a few nanometers [50, 51]. In our DFT calculations, we obtained the same qualitative results for both types of solvents. For brevity, we present the results for water with a reduced dielectric constant in the supplementary material (see Supporting Figure S9).

We first discuss the different geometrical arrangements considered for the Arg-containing complexes, which we show in Figure 5B: (i) T-shaped, (ii) coplanar stacked conformations, and (iii) dibridged hydrogen-bonded structures in the case of Arg-Tyr. These dimer structures are stabilized by different noncovalent interactions, characterized by noncovalent indices. Namely, we find wide areas corresponding to attractive van der Waals forces corresponding to π − π and CH-π interactions (represented in green in Fig. 5B) and dense attractive areas corresponding to hydrogen bonds (in blue). When comparing the values of ΔE_int_ for T-shaped and coplanar stacked configurations, our results indicate that the preferred geometry depends on the dielectric environment (see Figure 6A). At very low dielectric regimes, T-shaped structures are favoured in both Arg-Phe and Arg-Tyr complexes, whereas at higher dielectrics, stacked conformations predominate. This preference for the planar and H-bonded geometries is consistent with our observations from classical MD. In Supporting Figures S6 and S7, we show how the parallel and H-bonded configurations from DFT overlay with regions sampled during the MD simulations, while the T-shaped configuration is rarely found.

**FIG. 5.**
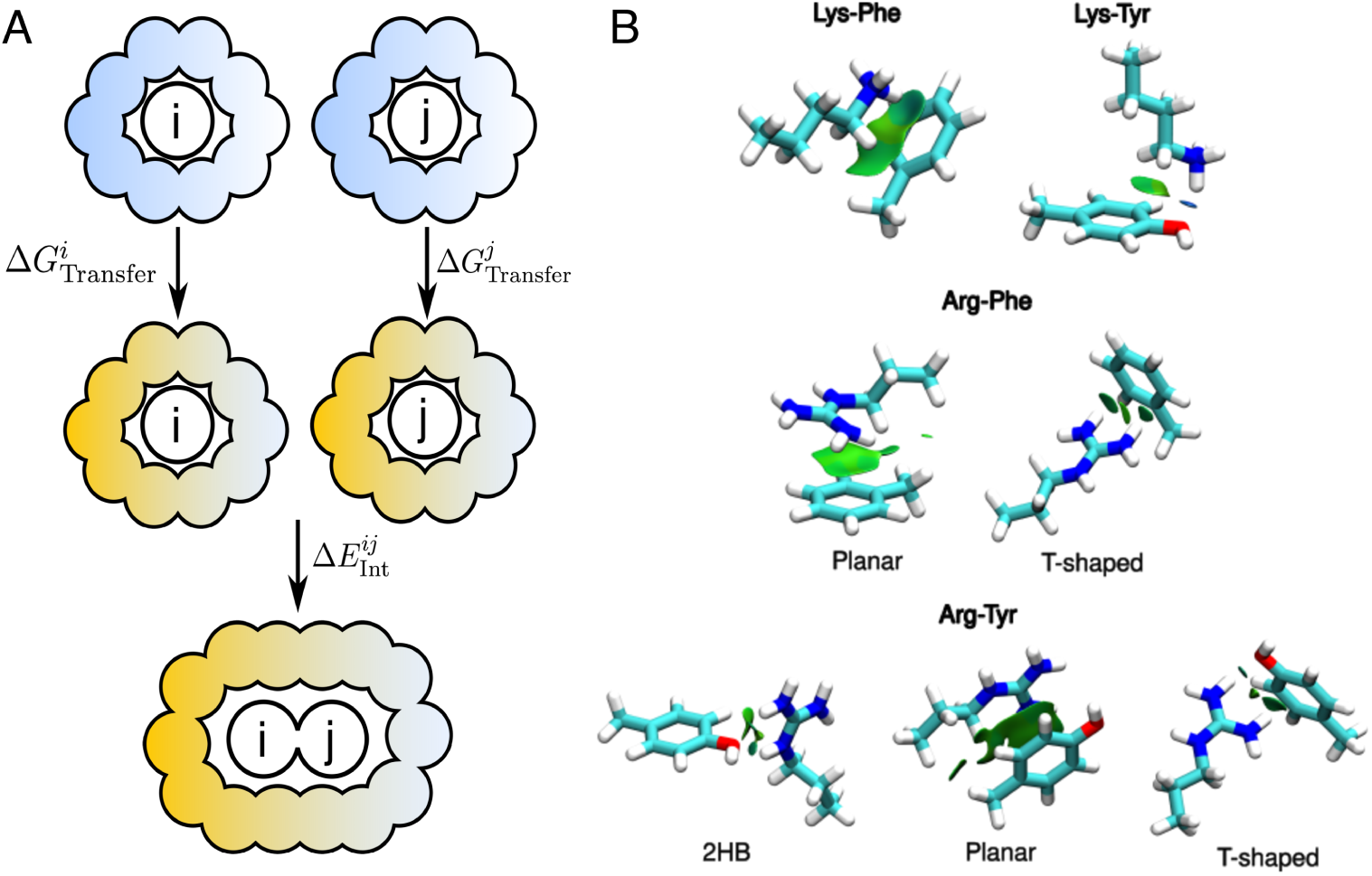
Contact formation energies from QM calculations. (A) Schematic of the process of contact formation for a pair of amino acid residues *i* and *j*. The first step corresponds to the transfer from water (blue), into a medium *s* (yellow), with an associated 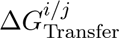. The second step involves the contact formation within the medium *s*, characterised with an energy 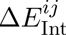. (B) Structures of all the dimers considered in the QM calculations. The surfaces shown are generated with NCI (Non-Covalent Interaction) analysis. In the case of Arg, various configurations were calculated: coplanar (P), T-shaped (T) and dibridged hydrogen bonded (2HB) in the case of the Arg-Tyr dimer.

**FIG. 6.**
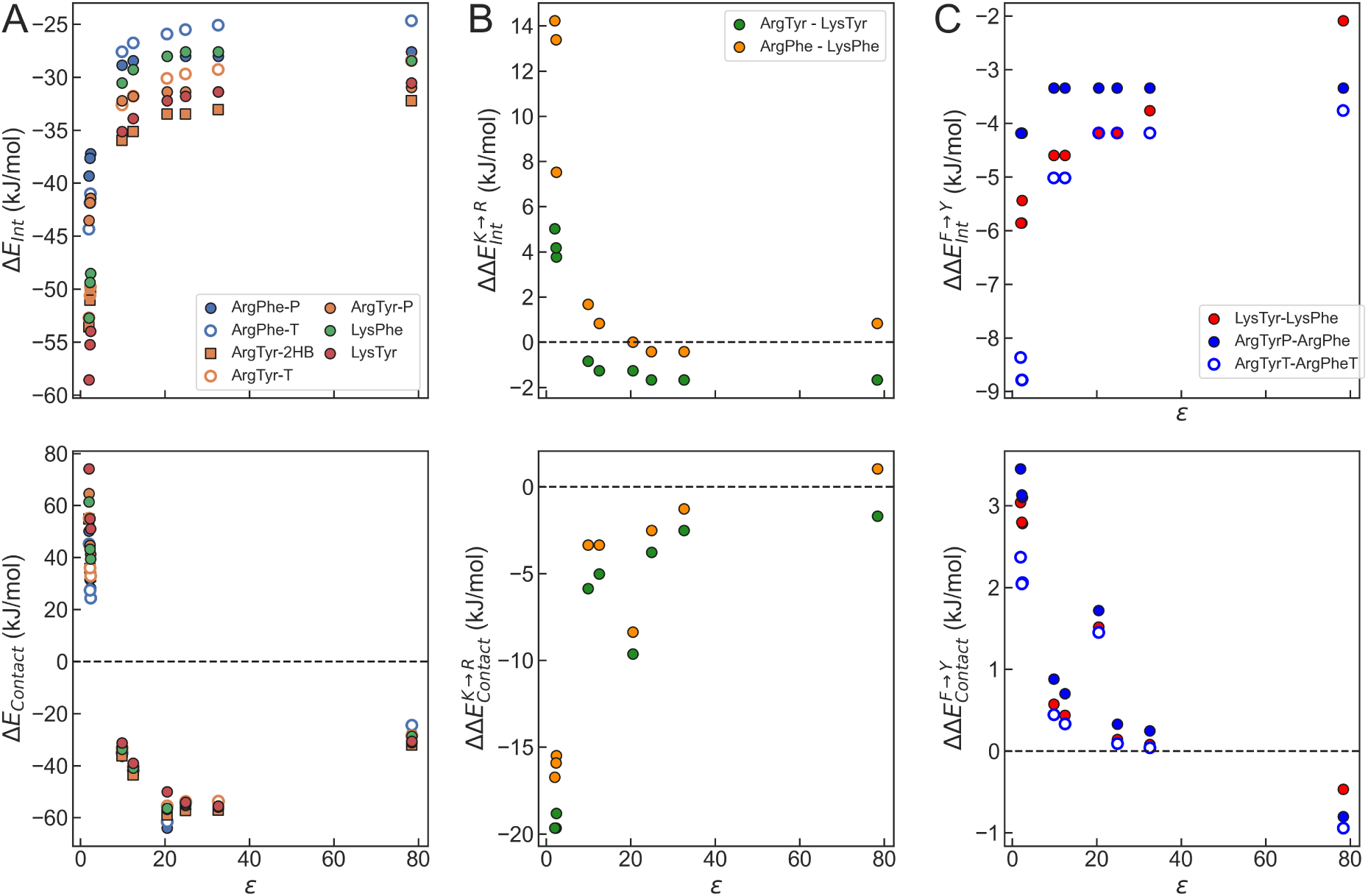
QM calculations for aromatic-charged dimers in different solvents. (A) Δ*E*_Int_ (top) and Δ*E*_Contact_ (bottom) energies for various configurations of the charged-aromatic residue dimers in various solvents as a function of the dielectric constant (*ε*). (B) ΔΔ*E*_Int_ and ΔΔ*E*_Contact_ energies for the Lys→Arg substitution in charged-aromatic dimers. We compare each Lys dimer with Phe and Tyr with the most stable conformation of the corresponding Arg pair at each dielectric value. (C) Same as B but for the Phe→Tyr substitution. ΔΔ*E* values were calculated by comparing geometrically analogous conformations for each pair.

The observed dependence with the solvent is consistent with earlier work by Chipot et al., who highlighted that Arg-containing cation-π complexes in the PDB predominantly adopt stacked geometries while perpendicular T-shaped motifs are favoured in the gas phase [52]. More recently, Calinski and Levy noted that although QM calculations support the stabilizing potential of π-stacked interactions, this need not necessarily translate into their prevalence in experimentally resolved protein structures [31]. This apparent discrepancy likely reflects the influence of the local environment, as interacting cation-π pairs are often situated in more hydrophobic regions rather than being solvent-exposed. The existence of a solvent effect is further supported by the positive correlation between the frequency of π-stacked conformations and the number of water molecules in the vicinity of the interacting pair. Consistent with this behaviour Cabaleiro-Lago et al. [53] showed that progressive microhydration of a guanidinium–aromatic dimer can shift the relative stability of different binding motifs from T-shaped to parallel conformations. They also reported that the interaction weakens systematically with increasing hydration, consistent with our observation that more polarizable environments reduce the magnitude of the interaction energy. Previous analyses show that π-stacked interactions are more frequent in solvent-exposed loops and turns [54] —and thus may be important stabilizers in disordered proteins— while protein interfaces also favour a coplanar arrangement of Arg’s guanidinium with aromatic rings [55]. Consistent with these observations, our results indicate that Arg–Tyr pairs prefer stacked cation–π conformations at medium and high dielectrics, whereas T-shaped geometries are more stable only at very low dielectric values.

In addition to cation-π interactions, hydrogen bonds have also been shown to be highly stabilising in these types of dimers, allowing Arg to form multiple interactions simultaneously with Tyr’s oxygen atom [31]. In our calculations, we find that the dibriged two hydrogen-bonded Arg-Tyr structure is the most stable Arg-Tyr dimer in a wide range of dielectrics. Although this propensity for hydrogen-bonded structures could be affected by the conformational restrictions of the peptide chain, in more than 60% of the cases analysed by Calinsky et al. of the PDB structures, the cation-π contact was accompanied by hydrogen bonds with Arg.

### Energetic analysis reveals the importance of the environment

So far, we have only considered interaction energies (ΔE_int_), representing the direct attraction between residues in a given environment. Their values reveal that the interaction strength between charged and aromatic residues is significantly amplified in low-dielectric environments (Figure 6A, top). Interestingly, while Arg-Tyr consistently shows the strongest interaction at higher and medium dielectrics, Lys-Tyr emerges as the preferred dimer at very low dielectrics. This aligns well with previous reports on the dielectric dependence of the interaction strength [56], and with the intuitive expectation that reduced dielectric screening enhances electrostatic forces.

However, the varying “stickiness” of distinct aminoacids requires not only the strength of their interactions inside condensates quantified above, but also their propensity to be transferred into the condensate from the dilute phase (see Figure 5A). Here, we estimate contact energies (ΔE_contact_) taking into account the cost of transferring the individual residues to that environment from the situation where they are fully solvated by water (i.e. ΔG_transfer_). The values of ΔG_transfer_ are quantitatively captured by the implicit solvation models we are using in our DFT calculations (see Methods and Figures S10 and S11). When we inspect the trends in ΔE_contact_ for the different cation-π pairs (Figure 6A, bottom), the picture fundamentally changes. Although contact formation remains favourable at intermediate and high dielectrics, at very low values of ε the contact energies become positive. This indicates that dimer formation is disfavoured in low dielectric organic solvents despite the strong attractions. This destabilisation is primarily driven by the unfavourable transfer energies of the charged residues from a high to a low dielectric environment. This highlights the importance of considering monomer transfer energies in the overall outcome of dimer contact energies, as they can dominate over the specific interaction energy of a dimer.

### Permutation of cationic and aromatic residues

To gain insight into the residue preference in cation-π interactions, we examined the relative interaction and contact energies after substituting Lys with Arg and Phe with Tyr (i.e. ΔΔE_int_ and ΔΔE_contact_, respectively, for K→R and F→Y). First we discuss the change in the cationic residue (see Figure 6B and supporting Figure S9). The value of 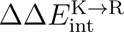 reveals an interesting crossover between Arg and Lys: at high dielectric values, Arg is preferred over Lys 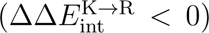 regardless of the aromatic partner. However, as the dielectric drops, this preference flips and Lys becomes favoured 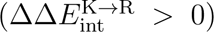. The presence of Tyr further enhances the preference for Arg at high dielectrics. Our findings are in agreement with the work of Kumar et al. [56], who reported that cation-π interaction strength diminishes with increasing values of ε, and Arg forms stronger interactions than Lys in water, despite being less favourable in the gas phase. They reasoned that while Lys-arene interactions are predominantly electrostatic and severely weakened by dielectric screening, dispersion and electrostatics are more balanced in Arg-arene pairs, making them far less sensitive to screening effects.

The picture for the contact energy difference between Arg and Lys pairs 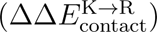 is strikingly different (see Figure 6B, bottom). Arg consistently remains the preferred charged residue throughout the dielectric range 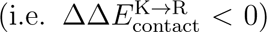, an effect that is particularly pronounced at low-dielectrics. This finding is explained by the more favourable transfer free energy of Arg compared to that of Lys. This provides a clear mechanistic explanation for the relevance of Arg as a sticker in biomolecular condensates. Its intrinsic interaction is more robust to dielectric changes, and its transfer to diverse environments is thermodynamically more favourable.

Our calculations also allow us to evaluate the effect of the F→Y substitution. In Figure 6C, we show the differences in interaction energies 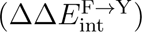 and contact energies 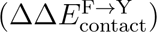) between dimers of the aromatics with either Lys or Arg. The interaction energy difference consistently indicates a general preference for Tyr in all dielectrics 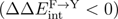 However, once again, trends in the contact energies are more subtle, showing a crossover in 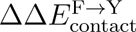 as the dielectric decreases. This finding mirrors our recent results on Phe and Tyr [17]. It is noteworthy that, although the general trends are similar when using the tuned water-dielectric model (Figure S9), the crossover appears at lower values of ε. This may be due to the capacity for the solvent to hydrogen bond with Tyr.

### Hydration free energy and sticker strengths in charged residues

The different trends between the interaction and contact energies reveal that the transfer-free energy is a key factor underlying the preference for Arg over Lys in cation–aromatic contacts. In Figure 7, we analyse this contribution separately, now considering both cationic and neutral forms of the side-chains of Lys and Arg. First, we show the solvation free energies of Lys and Arg (ΔG_Solv_, see Figure 7A), which is more negative for Lys than for Arg, reflecting the stronger localisation of the positive charge in Lys compared to Arg. When we instead consider the neutral forms of the amino acids, the trend reverses: Arg^0^ becomes better solvated than Lys^0^ in all solvents. These results are in agreement with experimental estimates of the hydration energy of charged residues and their neutral counterparts [57–59]. For the charged Lys/Arg side chains, the differences in solvation free energies are strongly attenuated in low-dielectric environments such as toluene, cyclohexane, and benzene.

**FIG. 7.**
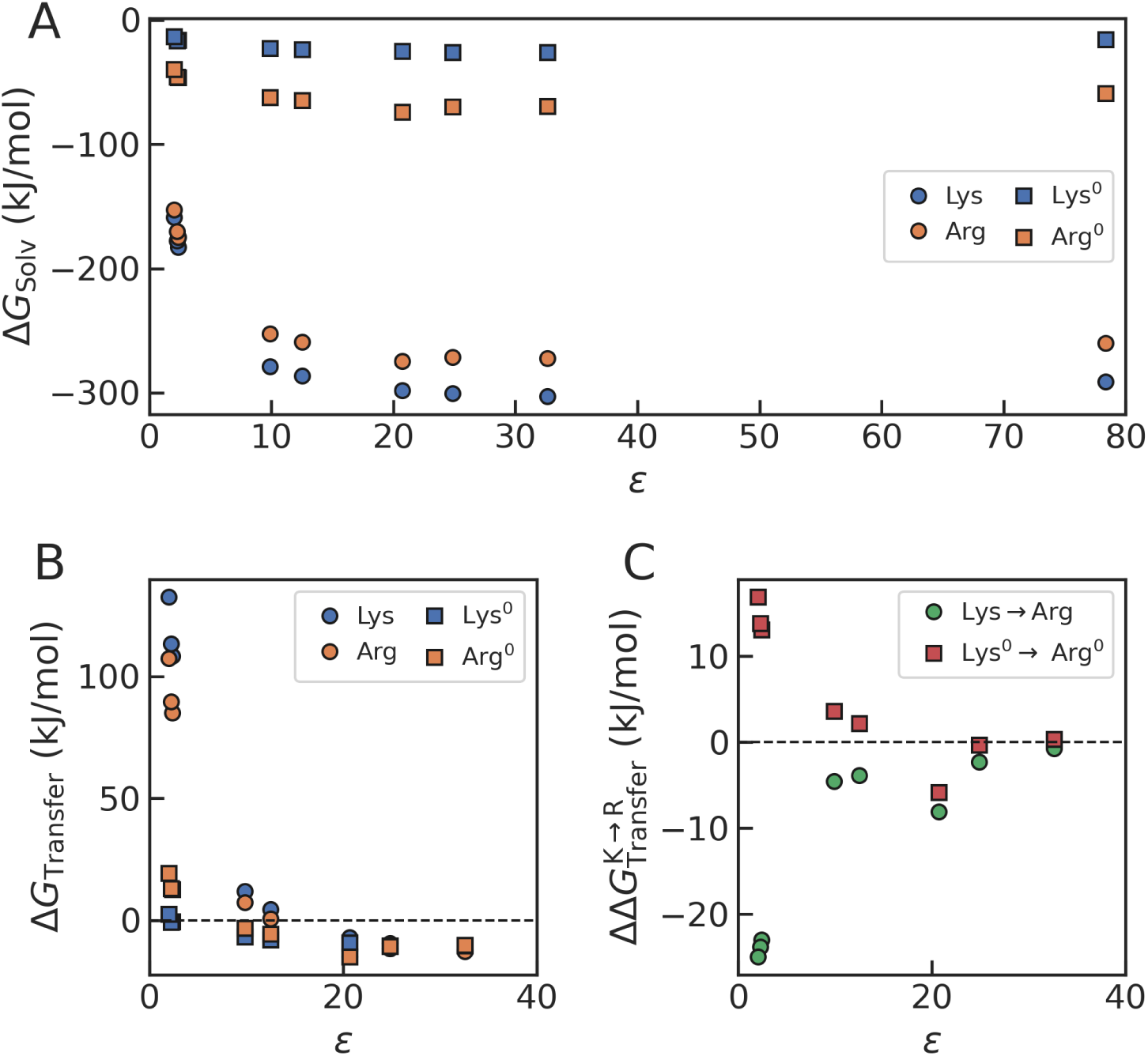
Solvation of charged and neutral amino acid side chain analogues of Lys and Arg. (A) Solvation free energy Δ*G*_Solv_ for charged and neutral Lys and Arg sidechain analogs at different solvents *s* as a function of their values of *ε*. (B) Transfer free energies (Δ*G*_Transfer_) estimated from the difference between solvation and hydration free energies (Δ*G*^solv^(*ε*) and Δ*G*^solv^(H_2_O)). (C) Transfer free energy differences 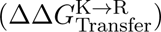 between between Arg and Lys.

From the solvation free energy in water and that of any other solvent s, we estimate the transfer free energy, i.e. ΔG_Transfer_(s) = ΔG_Solv_(s) − ΔG_Solv_(H_2_O), which we show in Figure 7B. At low dielectrics, the values of the transfer free energies are large and positive for both Arg and Lys due to the penalty of removing the positive charge from water. This penalty is consistently higher for Lys than for Arg. Hence, the relative transfer free energies 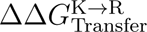 are negative, approaching –25 kJ/mol in low-dielectric solvents (see Figure 7C). In alcohols, the effect is less noticeable, with values of 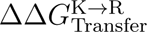 between –1 and –5 kJ/mol. These results are in semi-quantitative agreement with the trends observed in Fig. 3 for alcohols and acetone, and in qualitative agreement with those for aliphatic solvents. In contrast, when we consider the neutral form of the side chains of Lys and Arg, an opposite trend emerges. In this case, the energetic penalty for transfer from water to less polar solvents is much smaller, and Lys is now preferred over Arg. Accordingly, 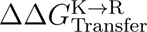 becomes positive and increases as the solvent polarity decreases.

The lower desolvation penalty of Arg relative to Lys is consistent with the results from MD simulations by Hong et al. [29]. From our simulations, we have estimated the radial distribution functions of water oxygen atoms around the central sidechain moiety (see Supporting Figure S8). In good accord with previous evidence, we find that, in the condensates, waters are slightly more likely to coordinate the amino group from Lys than the guanidinium in Arg. Together, these results indicate that the “stickier” nature of Arg compared to Lys arises from the difference between the very localized positive charge in Lys, which promotes hydration, compared to the delocalized positive charge of the guanidinium group, which has a lower penalty to transfer to nonpolar environments. Our findings confirm previous results [29, 60] that suggest that the differences in hydration free energy between Arg and Lys may explain their distinct roles as stickers. Our findings further indicate that charge neutralisation may play an important role in modulating the stickiness of amino acid residues.

## CONCLUSIONS

We have used a combination of classical MD simulations and quantum chemical calculations to study the relative strengths of Arg and Lys in biomolecular condensates. We have found that, contrary to what was observed for Tyr and Phe, cationic stickers preserve their hierarchy of interaction strengths —with Arg being stronger than Lys— at a wide range of values of the dielectric constant. In this sense, the hierarchy of sticker strength for charged residues is not context-dependent. However, there are subtleties when we try to understand the molecular determinants of this hierarchy. Much attention has been paid to the strength of cation-π interactions and the pseudoaromatic character of Arg [25, 28–31]. Our QM results show that the interactions become more favourable for Arg when the dielectric decreases, due to the decreased screening of the electrostatics in apolar media. But Lys interactions also become stronger, and even exceed the interaction strengths of Arg-arene pairs. The main contribution to the favorability of Arg over Lys comes from the large penalty of dehydrating the latter residue [29, 60].

Additionally, we have studied which of the aromatics makes a better partner with the cationic. We find that this depends on the effective permittivity of the medium, as described previously [17]. The greater sticker strength of Phe-Arg/Lys pairs relative to Tyr-Arg/Lys at low dielectrics is consistent with knowledge-based potentials derived from statistics of contacts in protein structures [18], where ε is known to be low [23].

Our work is necessarily limited due to the small size of the model peptides we are considering in our molecular simulations and the minimal representation of amino acid sidechains in the quantum chemical calculations. As we have explained before [17, 22], the computational convenience of short peptides comes at the expense of a high peptide density relative to that of true IDP condensates. We note, however, that our densities are consistent with those found for longer peptides by other authors [61–63] and the interaction patterns in peptide condensates agree with those of full length FUS and LAF-1 [64]. Additionally, the dielectric constant determined from the condensate simulations is in near-quantitative agreement with numerical estimates derived from experiment and simulations [24, 25]. All of this evidence suggests that peptide condensates are a good approximation to the condensates formed by disordered proteins.

In conclusion, our results provide a comprehensive description of cation-π interactions under varying dielectric conditions. We reveal that monomer transfer energies are often the dominant factor in determining residue preferences, leading to significant crossovers that defy predictions based solely on intrinsic interaction strength. This has profound implications for understanding the behavior of residues in confined biological environments, particularly within biomolecular condensates, where dielectric properties are altered. Our findings not only corroborate established biological observations regarding the prevalence of Arg-Tyr interactions and the importance of solvation but also offer novel mechanistic insights into how changes in the dielectric environment can fundamentally reshape the very forces governing protein-protein interactions and phase separation.

## METHODS

### Molecular simulations

#### Molecular models

We have run simulations of a short pentapeptide with sequence GGXGG with X=Arg or Lys capped at the N and C-termini in different media. All peptide structures were built using the LEaP program from Ambertools [65]. We used the Amber99SB⋆-ILDN force field, which includes corrections in backbone and sidechain torsions [66, 67] as provided within the PMX software for the generation of hybrid topologies [68]. The three-point TIP3P model was used for simulations including water molecules [69]. As alternative solvents, we use a collection of organic molecules with different dielectrics using the GAFF parameters from the Free Solvation Database by Mobley and co-workers [70].

#### Details of the simulations

We generated simulation boxes for the GGXGG peptide as before [17]. For simulations in pure water, we first inserted the fully extended structure of the peptide in an octahedral box, allowing at least 1 nm in each dimension. In the case of the organic solvents, we solvated the peptide using 500 molecules. Simulations of the peptide inside a condensate slab were performed by first inserting the GGXGG peptide in the center of a rectangular box of 10 nm×4 nm×4 nm and then surrounding it with a mixture of 300 copies of terminally capped Gly, Ser and either Phe or Tyr (which we term GSF and GSY condensates, respectively). This box was finally extended on both sides to reach a total length of 20 nm and filled with water molecules.

Due to the relevance of the electrostatic interactions of Arg/Lys to this study, we have tested different options for the neutralization of our simulation boxes. Whether ions are incorporated in the box can influence the results of the calculations, particularly when one considers non-homogeneous systems like our condensates in a slab geometry [71]. For this reason, we have carried out two batches of runs of condensate systems, one where a single Cl^−^ was added (termed “neutral”) and one where, in addition to this anion, we include ions up to 0.15 M salt (termed “physiological”). We have followed the same approach in the case of water. For other pure solvents, we rely on the uniform background charge added by the Ewald method.

We have equilibrated all our simulation boxes using a similar simulation protocol. First, solvated boxes were energy-minimized using a steepest descent algorithm. Second, we run a short simulation in the NVT ensemble using the Berendsen thermostat [72]. Next, we equilibrated the density of the simulation box in a short NPT simulation using the Berendsen barostat. Finally, we run long production simulations of up to 5 µs using the Parrinello-Rahman barostat [73] (see details in Table S1 in the Supporting Information). A leap-frog stochastic dynamics integrator with a 2 fs time step was used in the simulations. Van der Waals interactions were switched at 1 nm, and long-range electrostatics were calculated using the Particle-mesh Ewald method [74]. All the simulations were run using the Gromacs software package [75] (version 2024).

#### Analysis of the simulations

We have analysed the simulations using a combination of the Gromacs analysis programs [75], in-house Python scripts that exploit tools from the MDtraj library [76] and the scripts from the PMX package for the estimation of free energy associated with the alchemical transformation from the non-equilibrium runs [68, 77, 78]. These are derived from the work performed in the forward and backward non-equilibrium transitions, 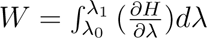. We report free energy estimates from Bennet’s Acceptance Ratio method [79] and refer the interested reader to the PMX papers for additional details [68, 77, 78]. Errors were determined via bootstrapping.

### Quantum chemical calculations

We use the Gaussian 16 program[80] to perform quantum mechanical calculations to study interaction energies of complexes formed by amino acid side chain analogues (see Figure 5B).

We consider all atoms starting from the C*^β^*, as in the past, Chipot and co-workers found that toluene was a better model than benzene for Phe sidechains in proteins [52]. We used Density Functional Theory (DFT) level, using the ωB97XD functional [81] and Pople’s 6-311++G(d,p) basis set [82] for geometry optimization. This methodology was succesfully used by Jorgensen et al. [47] to describe cation-π interactions. To visualize the nature of the non-covalent interactions in the optimized dimers, we performed a Non-Covalent Interaction (NCI) Index analysis [83]. To recapitulate the effects of different solvents, we run the geometry optimizations using the Polarizable Continuum Model approach [84]. In a given solvent, the interaction energy was calculated according to

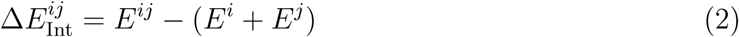

where i and j specify the interacting monomers. Notice that in continuum electrostatic models, the dielectric constant controls the energetic cost of solvating charged and polar groups: lower dielectric environments make the solvation of isolated ionic species less favorable, thus enhancing the relative stability of direct residue–residue interactions. Consequently, interaction energies depend on the dielectric properties of the surrounding medium even when the solvent is not explicitly present between the interacting residues. In addition, within self-consistent reaction field approaches, the polarizable medium can differentially polarize the electronic structure of the solute, providing a further dielectric-dependent contribution to interaction energies.

For a given monomer i, we calculate the free energy of transfer from water to a given solvent s using

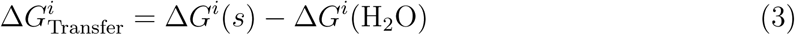

The free energies in both water and the solvent s are estimated using the SMD solvation model [85], using B3LYP/6-31+G(d,p) level of theory [86, 87]. As solvents, we have considered cyclohexane, benzene, toluene, n-octanol, 1-hexanol, acetone, methanol, and water. The adequacy of this approach in calculating transfer free energies has been tested with respect to experimental values of the solvation free energies in water and cyclohexane of neutral molecules analogs of aminoacid side chains, including n-butylamine and n-propylguanidine, sidechain analogs of Lys and Arg in their neutralized forms, in water and cyclohexane solvents. The corresponding SMD transfer free energies show a good agreement with the experiments (Fig S10 in supplementary material). In addition, SMD solvation free energies in water of a set of positively charged mimics relevant to Arg/Lys chemistry (guanidinium and alkyl-ammonium/alkyl-guanidinium series) were also compared with the experiments, finding very good agreement so that they further confirm the adequacy of our approach (Fig S11 in supplementary material).

On the other hand, we have used water as a solvent by varying the dielectric value, similar to our previous publication [17], in which this model was used in the context of aromatic-aromatic interactions. In continuum calculations, a reduced dielectric is used as an effective parameter to model water-like environments with a diminished collective polarization response (e.g. confinement, interfacial regions, and partial dehydration). This interpretation is consistent with recent work showing that the dielectric response of water can be strongly modified under confinement [88]. Combining the monomer transfer free energies with the dimer interaction energy at a given solvent s (see Figure 5A), we obtain the contact energy for the i-j pair:

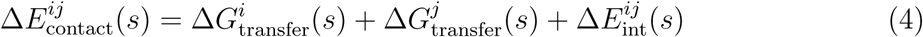

## ACKNOWLEDGMENTS

The work has been financed by the Basque Government through Project IT1584-22 and grant PID2024-158678NB-I00 funded by MICIU/AEI/10.13039/501100011033 and “ERDF A way of making Europe”. Lydia Armentia acknowledges support from the Jesús de Gangoiti Foundation. The authors also thank the IZO-SGI SGIker (UPV/EHU/ERDF,EU) DIPC for technical and human support and for the allocation of computational resources. The authors thankfully acknowledge RES resources provided by the Barcelona Supercomputing Center in MareNostrum5 (BCV-2025-2-0002).

