## Supporting information for "Arginine versus Lysine: Molecular Determinants of Cation–π Interactions in Biomolecular Condensates"

Table S1: Details of equilibrium MD runs for both $\lambda$ states. For each system, two independent equilibrium runs were performed corresponding to the $\lambda = 0$ and $\lambda = 1$ states. Box dimensions are values pre-equilibration. In the case of aqueous solvents, replicates of the simulations were run using both neutralised boxes and physiological salt concentration.

| System | Simulation time ($\mu$s) | # solvent molecules | # atoms | Box dimensions (nm/nm/nm) |
|---|---|---|---|---|
| $H_2O$ (neutral) | 2 | 2113 | 5590 | 5 / 5 / 5 |
| $H_2O$ (physiological) | 2 | 2103 | 5570 | 5 / 5 / 5 |
| Cyclohexane | 1.5 | 485 | 8796 | 5 / 5 / 5 |
| Benzene | 1.8 | 500 | 6066 | 5 / 5 / 5 |
| Toluene | 1.65 | 500 | 7566 | 5 / 5 / 5 |
| Octanol | 1.75 | 288 | 7842 | 5 / 5 / 5 |
| Hexanol | 1.93 | 379 | 8025 | 5 / 5 / 5 |
| Ethanol | 2 | 500 | 4566 | 5 / 5 / 5 |
| Methanol | 2 | 500 | 3066 | 5 / 5 / 5 |
| Acetone | 2 | 500 | 5066 | 5 / 5 / 5 |
| GSF (neutral) | 2 | 8140 | 43987 | 20 / 4.5 / 4.5 |
| GSF (physiological) | 2 | 8066 | 43839 | 20 / 4.5 / 4.5 |
| GSY (neutral) | 5 | 8035 | 43972 | 20 / 4.5 / 4.5 |
| GSY (physiological) | 5 | 7961 | 43824 | 20 / 4.5 / 4.5 |

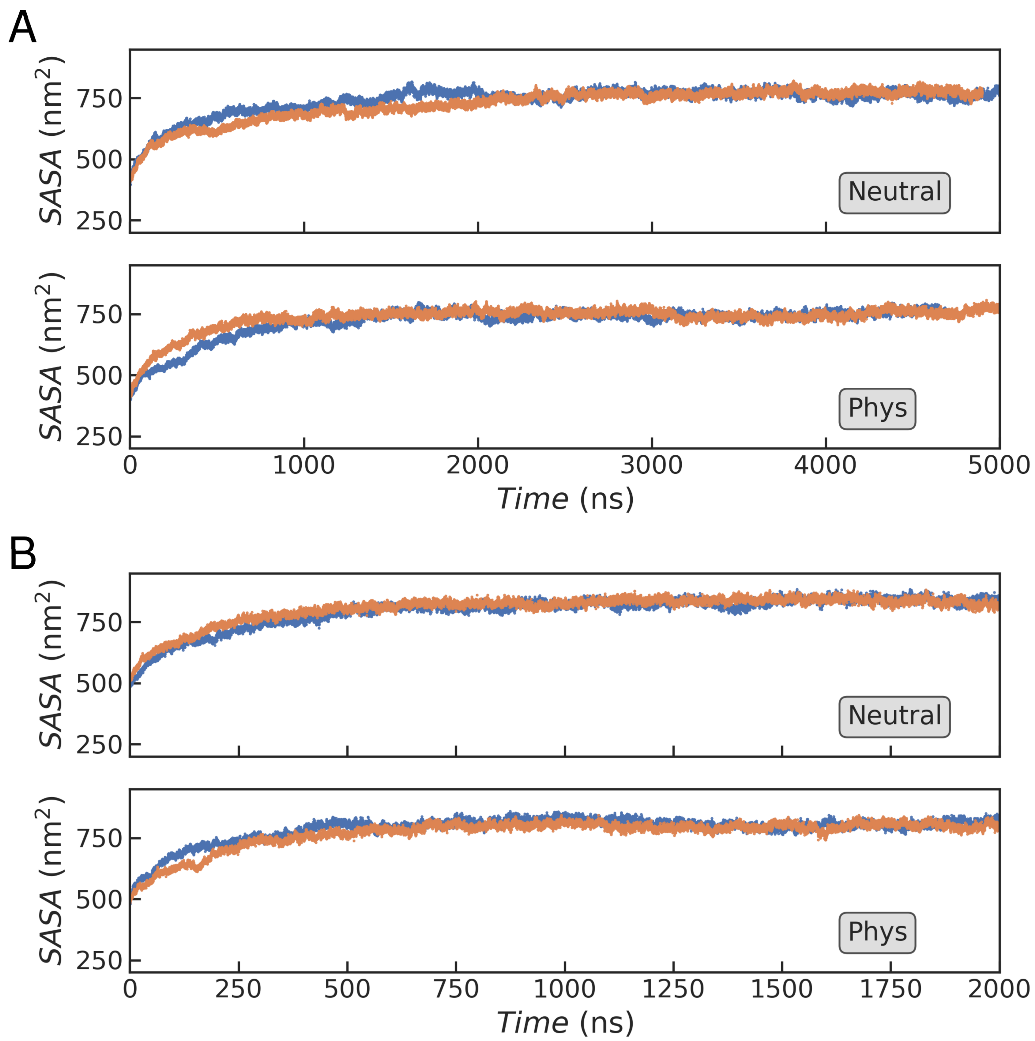


Figure S1: Solvent accessible surface area of all protein residues in the MD simulations of the GGXGG peptide within the GSY (A) and GSF (B) condensates. In each plot, the blue line corresponds to $\lambda = 0$ (GGKGG) and the orange line to $\lambda = 1$ (GGRGG).

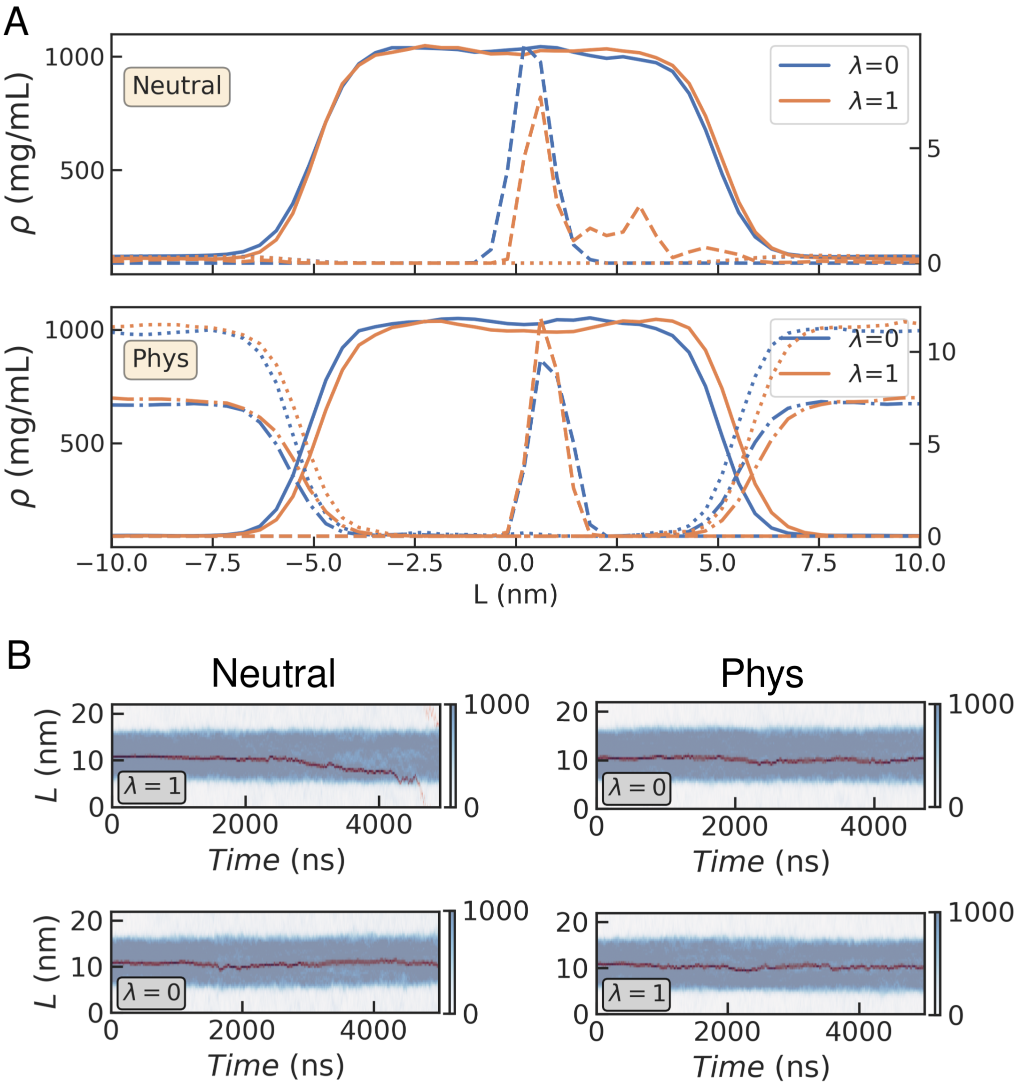


Figure S2: Density profiles for the GSY condensates. (A) Average over the simulation excluding the first 500 ns of trajectory for all residues (solid line), the K/R residue (dashed), $Cl^-$ ions (dashed) and $Na^+$ ions (dot-dashed). (B) Time series for all protein (blue) and K/R residue (dark red).

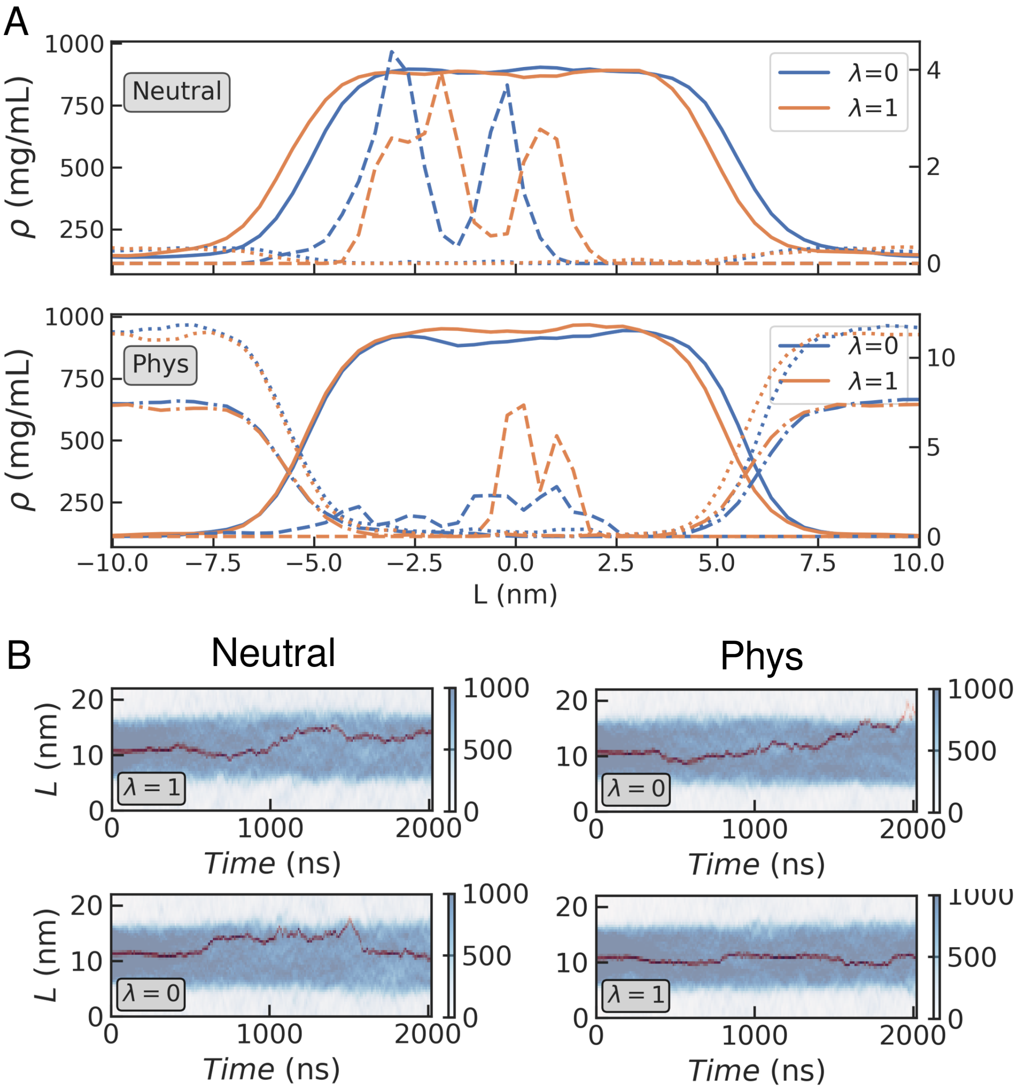


Figure S3: Density profiles for the GSF condensates. (A) Average over the simulation excluding the first 500 ns of trajectory for all residues (solid line), the K/R residue (dashed), $Cl^-$ ions (dashed) and $Na^+$ ions (dot-dashed). (B) Time series for all protein (blue) and K/R residue (dark red).

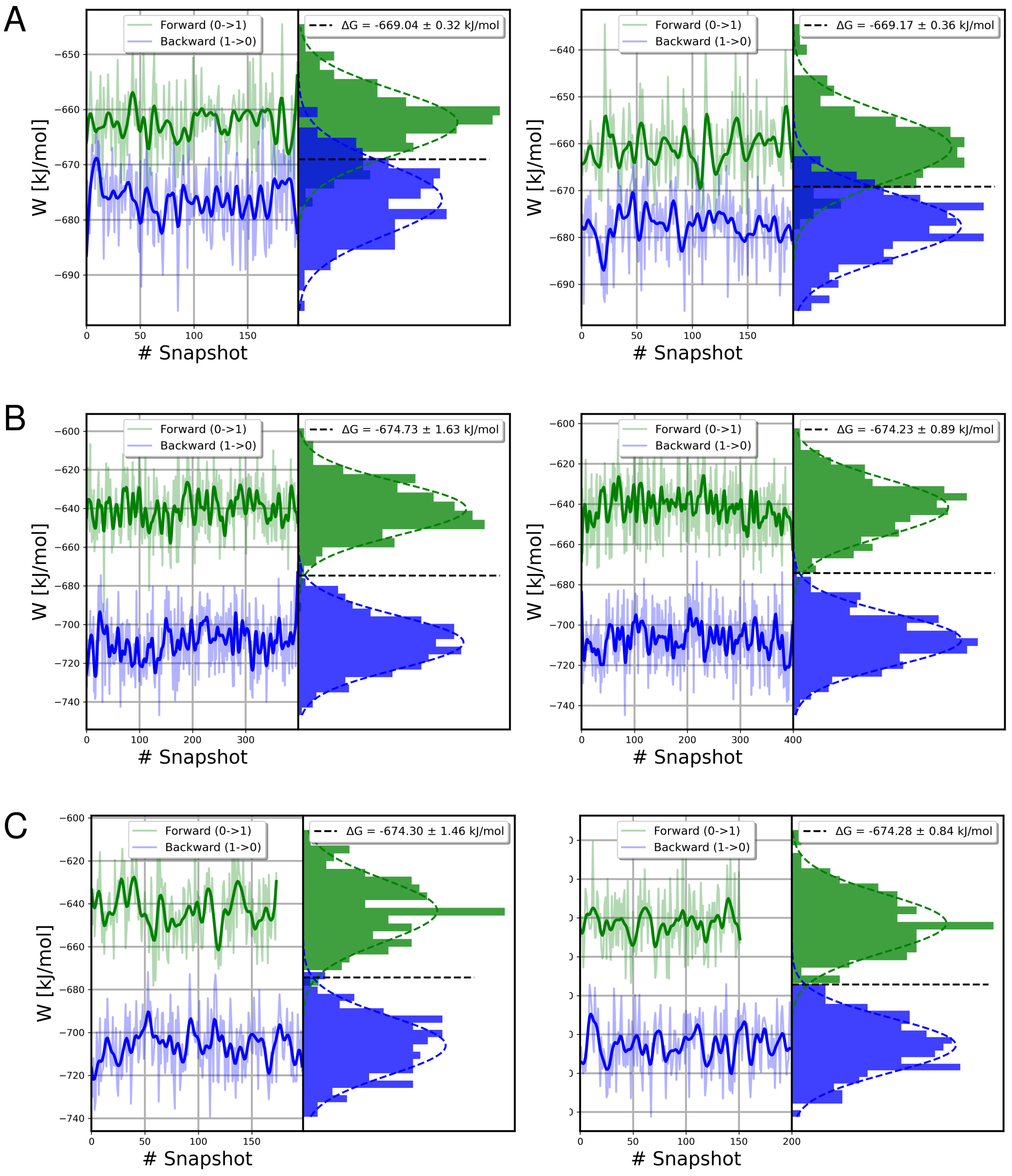


Figure S4: Time series for work for forward (0→1) and backward (1→0) alchemical transformations and corresponding distributions (left and right in each diagram, respectively). (A) Water, (B) GSY condensate and (C) GSF condensate. In all cases, we show the neutral system on the left and the physiological condition on the right.

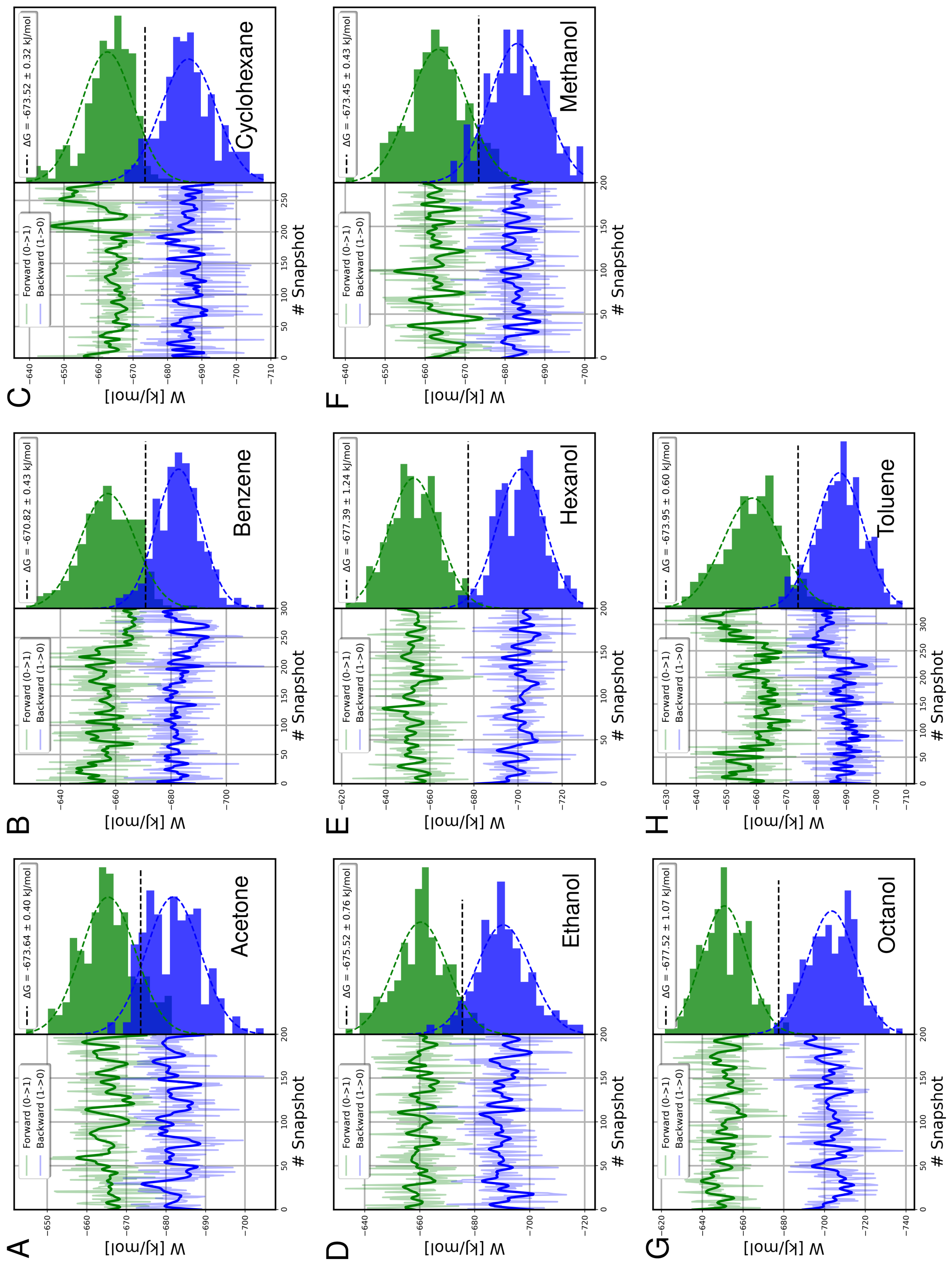


Figure S5: Work values for the forward and reverse alchemical transitions for different initial snapshots (left) and their distributions (right). The BAR estimator is shown in the legend. Each panel corresponds to a different solvent: (A) Acetone. (B) Benzene. (C) Cyclohexane. (D) Ethanol. (E) Hexanol. (F) Methanol. (G) Octanol. (H) Toluene.

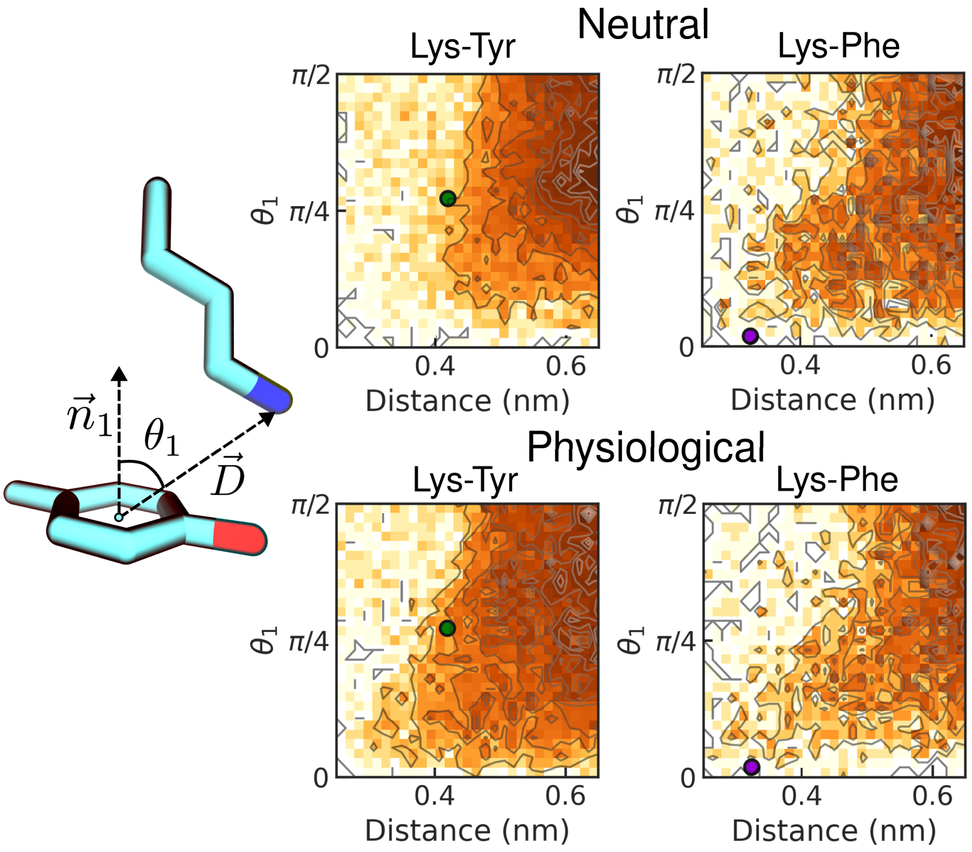


Figure S6: Statistical analysis of the interactions between Lys and aromatic residues in the MD simulations (contours). In the diagram on the left, we define the selected geometrical parameters:[S1] the distance of the vector $\vec{D}$ between the NZ atom and the ring centroid and the $\theta_1$ angle between $\vec{D}$ and the vector $\vec{n}_1$ normal to the ring plane. We show results at both neutral (top) and physiological (bottom) conditions for the simulations in the GSY (Lys-Tyr) and GSF (Lys-Phe) condensates. Green and purple circles correspond to the values of the same geometrical parameters from the Lys-aromatic structures from the QM calculations in water.

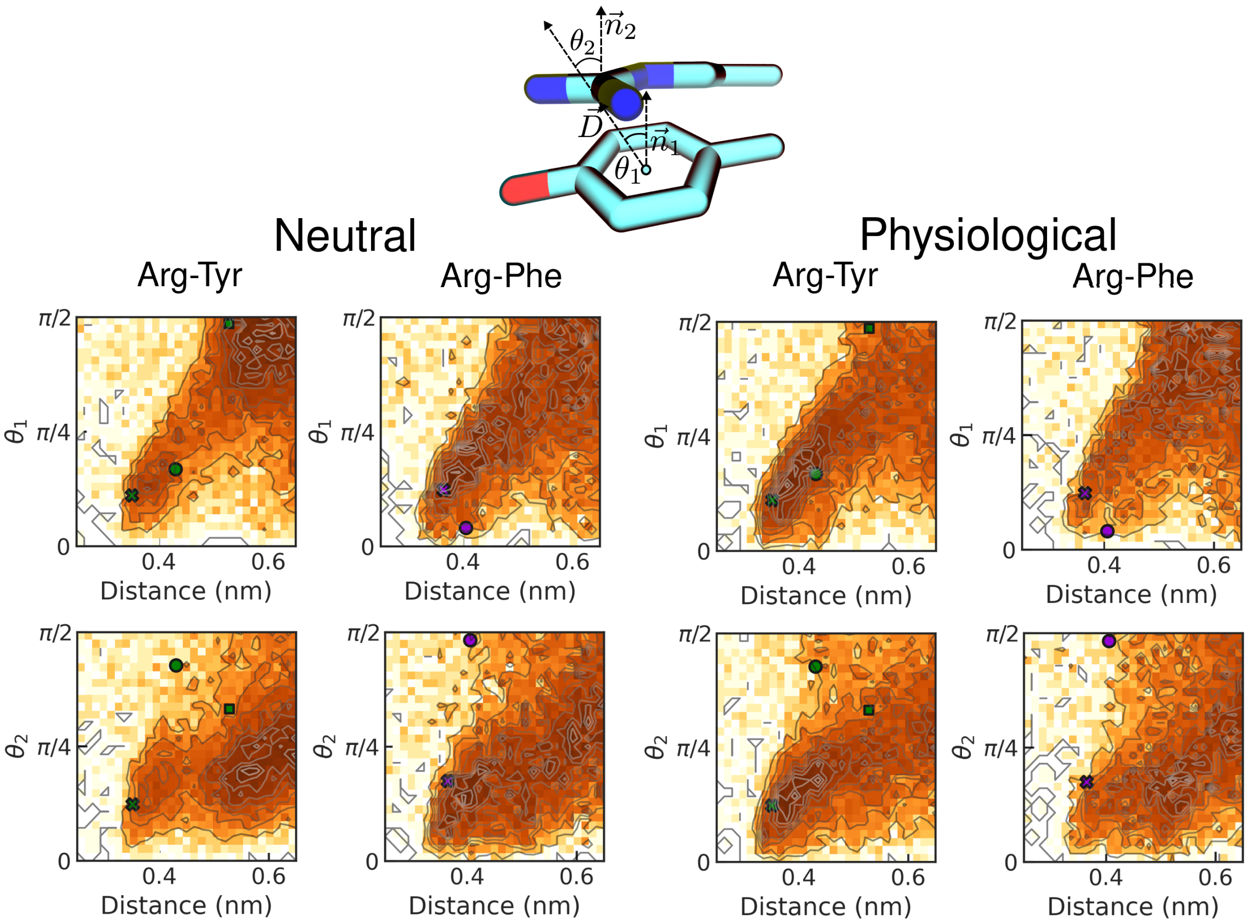


Figure S7: Statistical analysis of the interactions between Arg and aromatic residues in the MD simulations (contours). In the diagram on the top, we define the selected geometrical parameters:[S1] the distance of the vector $\vec{D}$ between the center of the guanidinium group and the ring centroid, the $\theta_1$ angle between $\vec{D}$ and the vector $\vec{n}_1$ normal to the ring plane, and the $\theta_2$ angle between $\vec{n}_1$ and the vector $\vec{n}_2$ normal to the guanidinium plane. We show results at both neutral (left) and physiological (right) conditions for the simulations in the GSY (Arg-Tyr) and GSF (Arg-Phe) condensates. Green and purple circles correspond to the values of the same geometrical parameters from the Arg-aromatic structures from the QM calculations in water. Stars mark the parallel configuration, circles the T-shaped, and in the case of Arg-Tyr, squares mark the H-bonded.

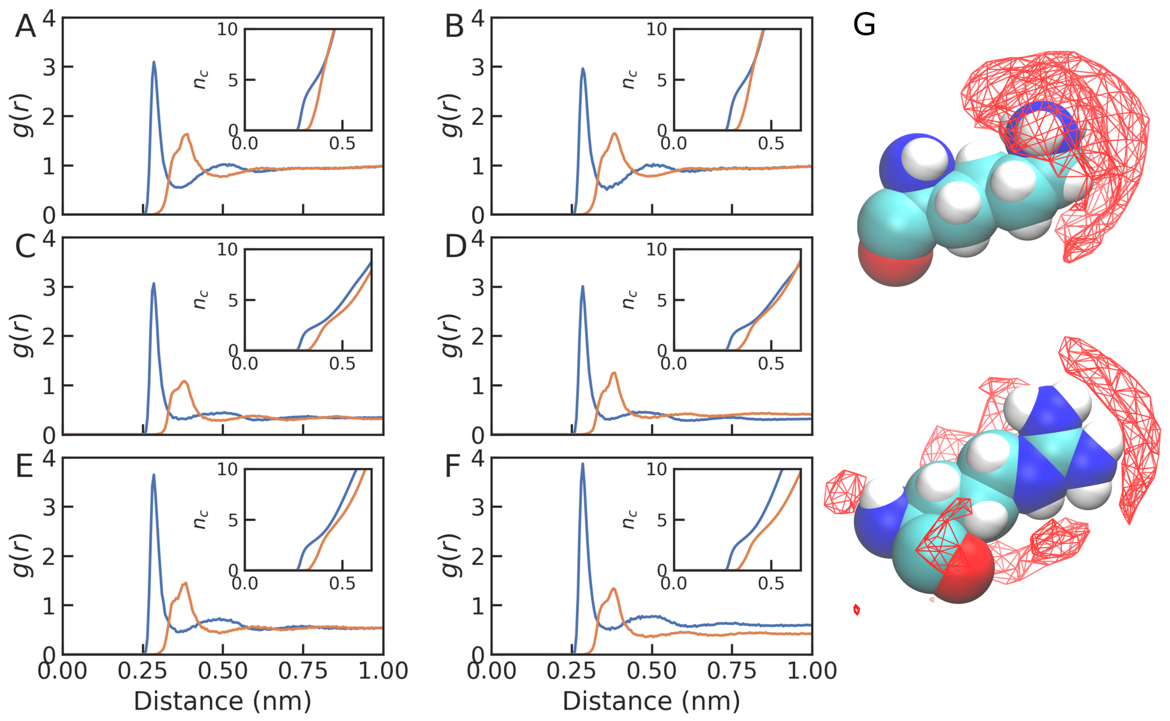


Figure S8: Radial distribution function for the distance between the central atom of the amonium or guanidinium groups and the water oxygen in different simulation datasets. (A, B) water; (C, D) GSY; (E, F) GSF. (A,C,E) neutral and (B, E, F) physiological. (G) Cartoon representation of the spatial distribution of water oxygens around the central atom in Lys (top) and Arg (bottom).

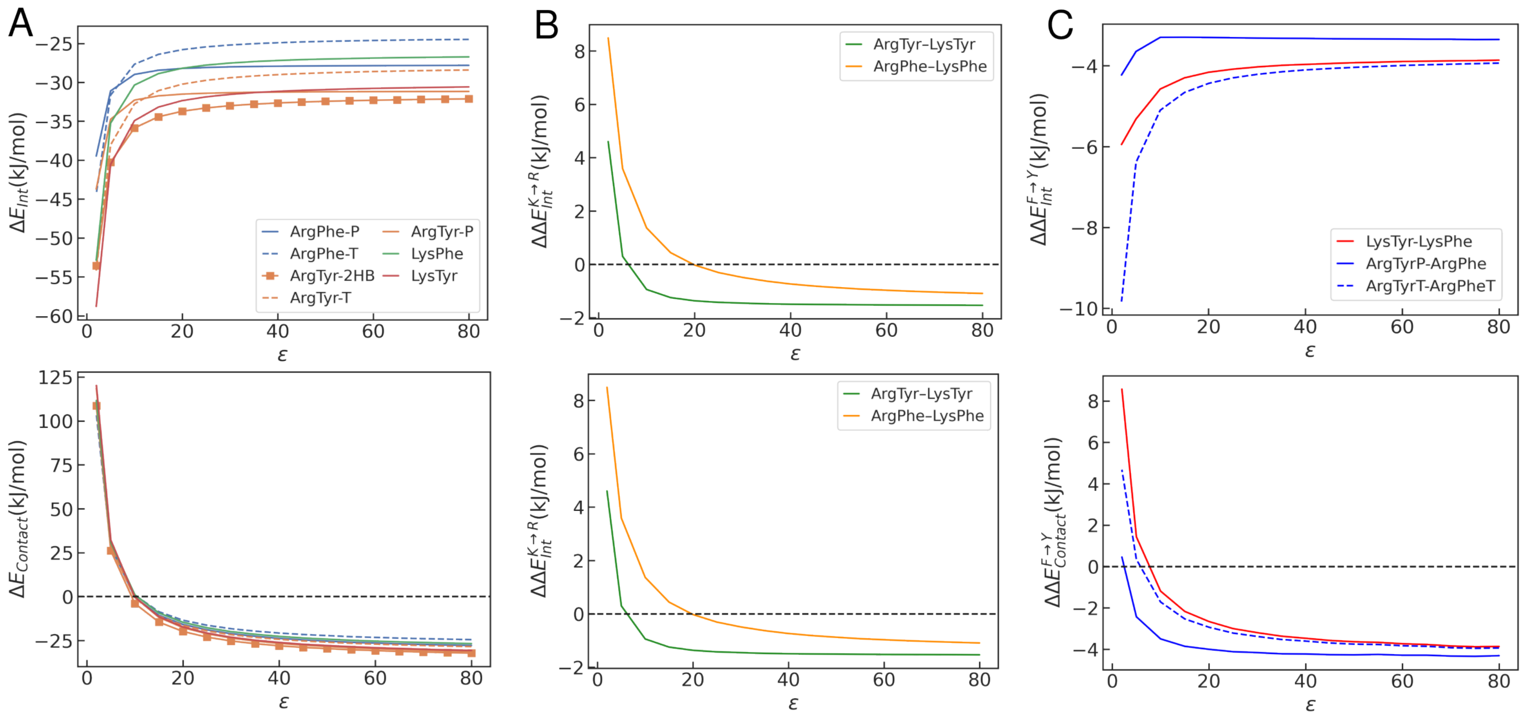


Figure S9: (A) $\Delta E_{\mathrm{Int}}$ (top) and $\Delta E_{\mathrm{Contact}}$ (bottom) energies for various configurations of the charged-aromatic residue dimers in various solvents as a function of the dielectric constant ($\varepsilon$). (B) $\Delta\Delta E_{\mathrm{Int}}$ (top) and $\Delta\Delta G_{\mathrm{Contact}}$ energies for the Lys→Arg substitution in charged-aromatic dimers. We compare each Lys dimer (Lys-Phe, Lys-Tyr) with the most stable conformation of the Arg pair (i.e. Arg-Phe, Arg-Tyr) at each dielectric value. (C) Same as B but for the Phe→ substitution. The energy differences were calculated by comparing geometrically analogous conformations for each pair.

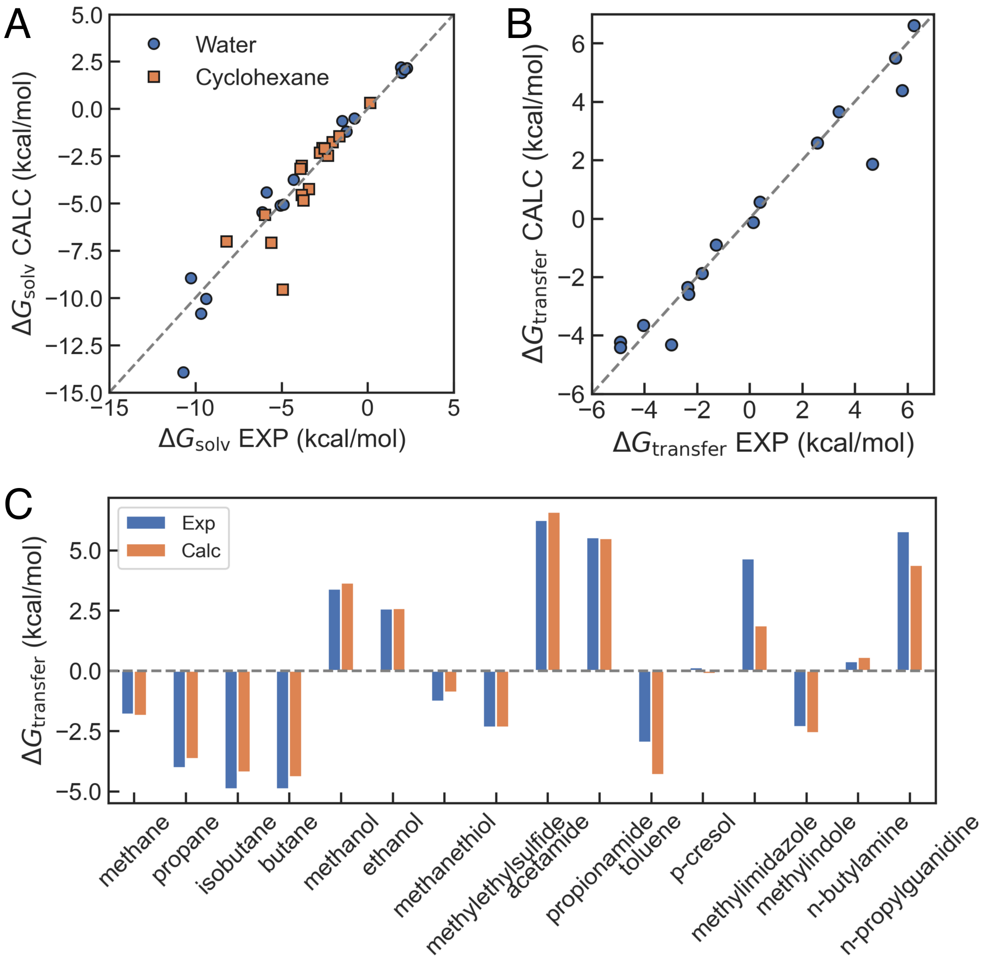


Figure S10: Comparison between DFT calculations on sidechain analogues using SMD and experiments. (A) Correlation between experimental and calculated solvation free energies in water and cyclohexane. (B) Correlation between experimental and calculated transfer free energies. (C) Bar chart of transfer free energies. Experimental data was taken from Chang et al.[S2]

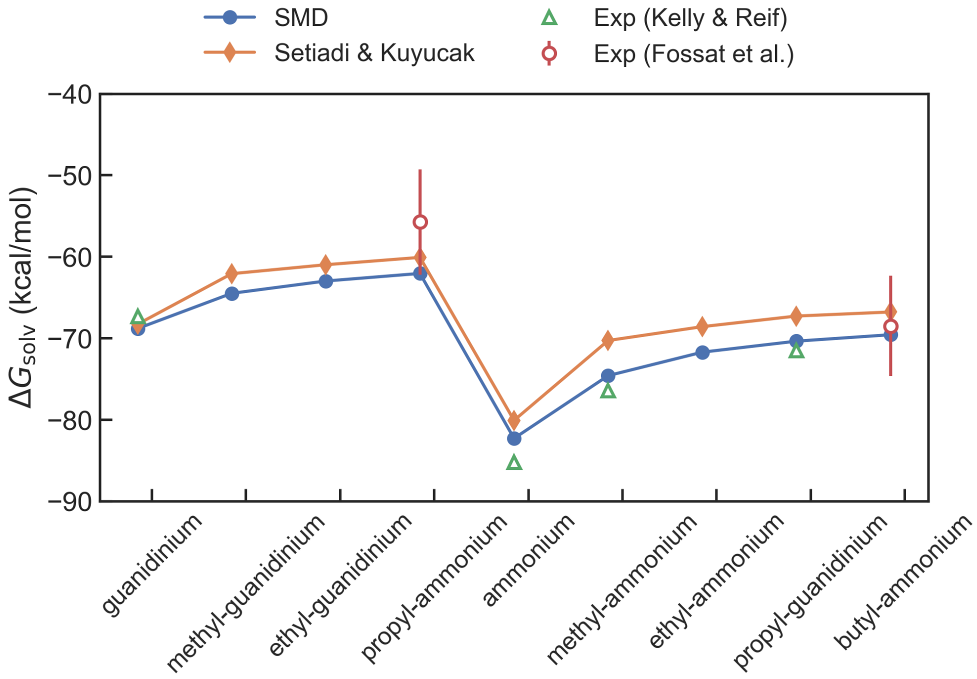


Figure S11: Solvation free energies on guanidinium and ammonioum variants calculated using SMD. We compare against previous calculations[S3] and experimental estimates.[S4–S6]
